# Microbial transformation of traditional fermented fertilizer bokashi alters chemical composition and improves plant growth

**DOI:** 10.1101/2021.08.01.454634

**Authors:** Nisreen Abo-Sido, John W. Goss, Alden B. Griffith, Vanja Klepac-Ceraj

## Abstract

Bokashi is an organic soil amendment that makes use of microbial processes to break down agricultural waste and create a nutrient-rich fertilizer. The benefits of various types of bokashi on soil fertility and plant growth are well documented, however the changes in microbial community composition and nutrients during bokashi maturation remain poorly characterized. Here, we aimed to identify potential differences in the quality of bokashi made using different ingredients and to investigate the biochemical transformation and microbial community succession of bokashi throughout the maturation process. We compared the effects of these different types of bokashi on the growth of cucumber (*Cucumis sativus*) and kale (*Brassica napus subsp. pabularia*) seedlings, measured concentrations of NH_4_^+^ and PO_4_^3-^, and characterized the bokashi bacterial and fungal communities over a 12-day maturation period. We found that cucumber and kale plants growing in all types of bokashi-amended soils exhibited increased chlorophyll levels and dry biomass. During bokashi maturation, we observed a decrease in available PO_4_^3-^, and an increase in NH_4_^+^. There also appeared to be an increase in relative abundances of decomposers and beneficial microbes and a decrease in putative plant pathogens. Regardless of starting bokashi ingredients and differences in microbial composition and nutrient trends, all types of bokashi similarly improve plant growth and contain beneficial microbes.

## 1. Introduction

Beginning in the 1960s, the Green Revolution marked an era of high-yield agriculture, made possible by the Haber-Bosch process, which converts atmospheric nitrogen into ammonia for fertilizer. Presently about half of the world population relies on this increase in yield for sustenance (Erisman et al., 2008). However, it has become increasingly evident that this boom in agricultural yield is unsustainable: depending on factors such as climate, soil type, and application technique, half or more of applied fertilizers are lost to the environment (Good and Beatty, 2011; Sutton et al., 2011; Weisler et al., 2001), polluting bodies of water, causing eutrophication, emitting greenhouse gases, and presenting human health risks (Erisman et al., 2013; Fowler et al., 2013). Furthermore, dependence on chemical agriculture has debased farmer knowledge. Mediating these negative externalities is very costly (Sutton et al., 2011), as is the cost of fertilizer to farmers. Still, supplying sufficient nutrients is critical to meeting yield demands. Amending soils using agroecological approaches may optimize natural nutrient cycling processes and effectively limit the negative consequences of high fertilizer inputs.

Many small-farmer agroecological techniques can meet high yields and foster biodiversity while promoting dynamic nutrient cycling (De Schutter, 2011; Kremen, 2015). Organic amendments can improve soil composition by increasing concentrations of phosphorus, organic carbon, and nitrogen pools. Global reports argue that small farmers can double food production by utilizing these already existing agroecological methods, such as polycultures, agroforestry, cover-cropping, and many others (Altieri et al., 2012; De Schutter, 2011; IAASTD, 2009). An example of such a method is a traditional East-Asian fertilizer called *bokashi*, which is an organic soil amendment that makes use of microbial processes to break down agricultural waste and create a nutrient-rich fertilizer.

Bokashi can be made with variable combinations of resources, depending on what is available regionally and financially, so long as some source of nitrogen, carbohydrate, and active microbes are included and processes foster partially anaerobic conditions. This makes bokashi far more affordable and accessible than commercial fertilizers. Unlike compost, bokashi matures rapidly, in just two weeks, and fundamentally integrates animal waste. Various studies have illustrated the efficacy of bokashi as a fertilizer that improves plant growth (Álvarez-Solís et al., 2016; Aurora Gomez-Velasco et al., 2014; Bautista-Cruz et al., 2014; Boechat et al., 2013; França et al., 2016; Jaramillo-López et al., 2015; Lima et al., 2015; Peralta-Antonio et al., 2014) and several have investigated the amendment’s nutrient content (Álvarez-Solís et al., 2016; Aurora Gomez-Velasco et al., 2014; Boechat et al., 2013; Lima et al., 2015; Peralta-Antonio et al., 2014).

Though the benefits of various types of bokashi fertilizer on soil fertility and plant growth have been explored, few studies exist that compare the effects of different ingredients on bokashi efficacy (Boechat et al., 2013; Lima et al., 2015). Even fewer analyze the biochemical transformations of bokashi from starting ingredients and the mechanisms underlying the fertilizer’s effectiveness (Boechat et al., 2013). Moreover, researchers have identified possible microbes present in bokashi (Magrini et al., 2011; Yamada and Xu, 2001), but the relative microbial community composition of bokashi throughout the maturation process has not been described. Understanding nutrient transformation in conjunction with microbial community composition in the maturation of bokashi could provide mechanistic insight not only into the decomposition processes within bokashi piles, but also the efficacy of bokashi in supplying plant nutrients. In effect, an understanding of these processes could inform methods for optimizing bokashi production from available waste resources and contribute to the adaptability of the fertilizer to different contexts.

The objectives of this study were to both identify potential variation in bokashi quality made using different ingredients and to investigate the microbial and chemical transformation of bokashi during maturation. We posed the questions: (1) Are different types of bokashi more or less effective in improving plant growth? (2) How does the resulting nutrient composition of bokashi differ if different starting ingredients were used? (3) Which microbes convert bokashi ingredients into an effective fertilizer? To examine these questions, we measured the nutrient composition and characterized the microbial community associated with maturing bokashi made with different starting materials. Furthermore, we compared the effects of these different types of bokashi on the growth of cucumber (*Cucumis sativus*) and kale (*Brassica napus subsp. pabularia*) seedlings. Here we report data on the ammonium (NH_4_^+^) and phosphate (PO_4_^3-^) concentrations and the bacterial and fungal makeup of bokashi over a 12-day maturation period.

## 2. Materials and methods

### 2.1. Making bokashi and collecting samples

We used three different recipes for making bokashi across two experiments (Figure 1; Exp. 2 began five days after the completion of Exp. 1). All bokashi types included 2 gallons of cow manure (collected from the Natick Community Organic Farm, Natick, MA, USA), 3 gallons of soil (“Coast of Maine Topsoil” made from loam, compost, and peat), and 1 gallon of corn flour (Bob’s Red Mill organic whole grain corn flour). In Experiment 1, both bokashi types received 5 g *Saccharomyces cerevisiae* (Fleischmann’s^®^ Active Dry yeast) and 50 mL molasses (Crosby’s Fancy Molasses). The estimated number of yeast cells per gram of starting bokashi mixture is 3 x 10^6^. The control (“Exp. 1 - Control”) included 1 gallon of raw rice hulls while the manipulated type (“Exp. 1 - Charcoal”) received 1 gallon of rice hull charcoal. Rice hulls were smoked to charcoal for 4 hours. In Experiment 2, both bokashi types were made with 1 gallon of raw rice hulls, but the control (“Exp. 2 - Control”) was made with 5 g *S. cerevisiae* and 50 mL molasses, while the manipulated type (“Exp. 2 - IMO”) was made using IMOs with 50 mL of molasses. IMOs were collected from visible microbial growth on rice that had been covered in local soil for three days and then propagated in a sealed jar with brown sugar at a ratio of 1 part sugar to 3 parts rice IMOs. After 3 days in the jar, the IMO mixture was added to a pile of 1 gallon each of soil and corn flour, and some water, then covered and mixed daily for 1 week. IMOs were applied as a bokashi ingredient by adding 2 handfuls of this IMO-soil to the bokashi pile. Note that the controls in both experiments were made using the same procedure and ingredients, but the age of the manure differed for the two experiments.

**Figure 1:**
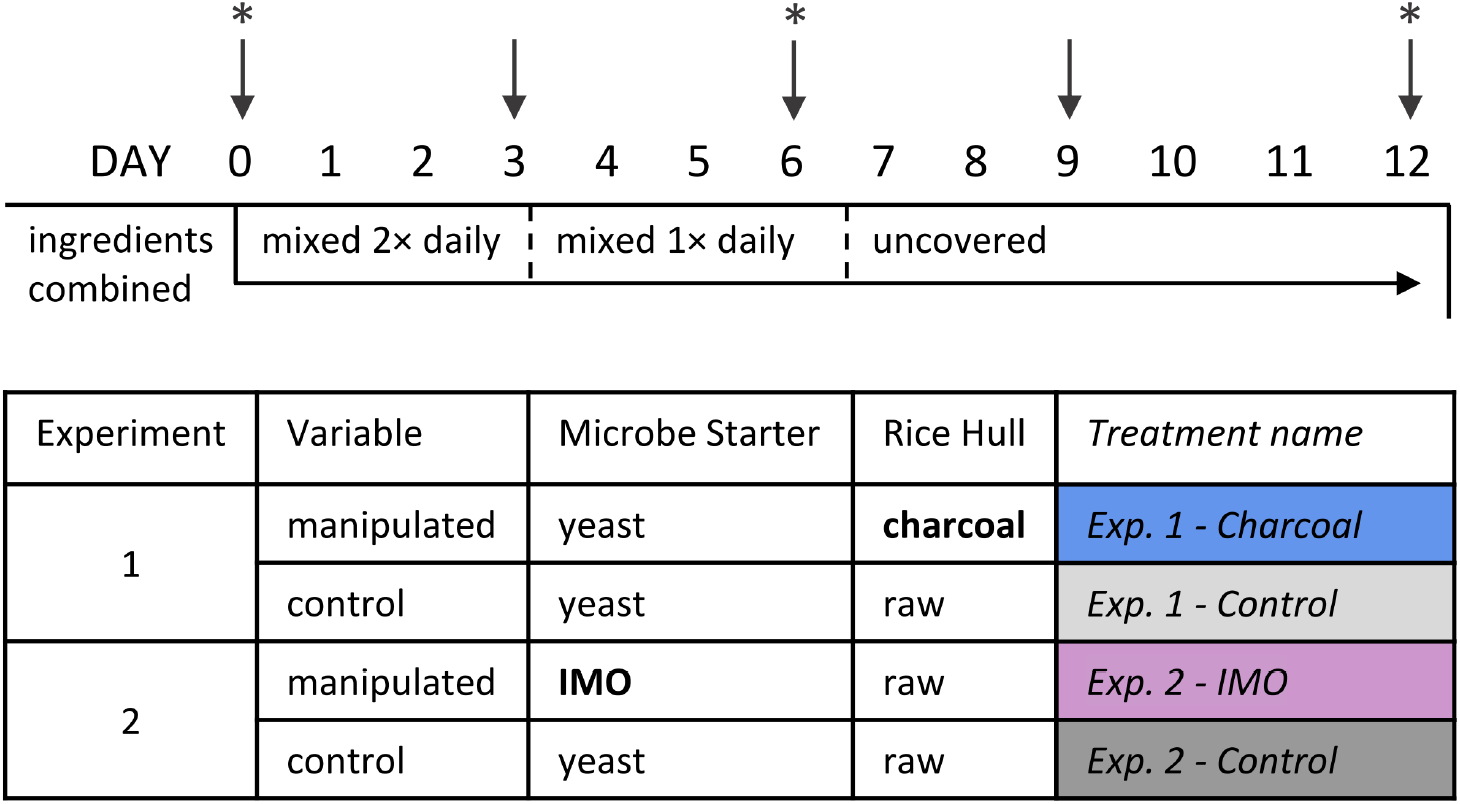
Method for making bokashi. Arrows indicate sampling days for nutrient and 16S rRNA gene amplicon analyses; stars indicate samples taken for ITS amplicon analysis. Table describes differences in composition of bokashi types.

Bokashi piles were made in two rounds of experiments inside the same greenhouse, separated only by time, so the manure for the bokashi was collected at different times. The temperature of the greenhouse was maintained at approximately 22-25 °C, and four replicates of each variation of bokashi were made in separate piles. All the ingredients were placed on black plastic tarps on tables in the greenhouse and water was added to moisten the piles to approximately 60%, amounting to 1.5 L of water per pile. The piles were covered with tarps for the first seven days and mixed twice daily for the first three days, then once daily for the following 8 days, until fully dry on day 12 (Figure 1). Samples were collected on days 0, 3, 6, 9, and 12 for the nutrient and bacterial (16S rRNA gene amplicons) analyses; samples collected on days 0, 6, and 12 were processed for fungal (ITS amplicons). All samples collected for 16S and ITS analyses were immediately frozen at −20 °C until further processing.

### 2.2. Plant growth experiments

Kale (*Brassica napus subsp. pabularia* - Red Russian) and cucumber (*Cucumis sativus* - Green Finger) seedlings were treated with the four bokashi treatments (Exp. 1 - Charcoal, Exp. 1 - Control, Exp. 2 - IMO, Exp. 2 - Control), compost, or the control condition of soil alone. The compost was Miracle Grow’s “Nature’s Care: Really Good Compost.” The base soil was the same soil used to make bokashi.

Kale and cucumber seedlings were started in plug trays with seed starting mix. Approximately three weeks after germination, the cucumber and kale seedlings were transplanted into 400-mL and 500-mL pots, respectively, with one seedling per pot. To replicate farmer methods, each pot contained one handful of either bokashi (60 grams), compost (80 grams), or soil (140 grams), between two handfuls of soil. In effect, we created a soil-barrier between the amendments and the seedlings to avoid nutrient burn (Rivera, 2001). Each of the 6 treatments were replicated 8 times per species, totaling 96 pots: 2 species × 6 treatments × 8 replicates. Within the 8 replicate pots, each of the 4 original bokashi replicate piles were represented twice. Plants were watered as needed and received ambient light.

Three weeks after seeds were sown, we estimated the chlorophyll concentration of two mature leaves per plant using the Opti-Sciences CCM-300 Chlorophyll Content Meter (values are expressed as the fluorescence ratio of 735 nm / 700 nm; Gitelson et al., 1999). Then we harvested the aboveground tissue and dried them in a 65 °C oven for approximately 2 days before measuring dry biomass.

### 2.3. Preparing samples for discrete nutrient analysis

Collected bokashi samples were dried (45 °C for 2 days) and ground for 2 min via a mixer mill, and the powdered samples were used to make extracts for colorimetric determination of NH_4_^+^ (ammonium), NO_3_^-^ (nitrate), and PO_4_^3-^ (phosphate) by a Discrete Analyzer (Astoria-Pacific, Inc., Clackamas, OR, USA). Test extracts for NH_4_^+^and NO_3_^-^ were made by combining 7 g of sample and 40 mL of 2 M KCl to a 50-mL centrifuge tube, shaking the samples for an hour, allowing samples to settle for an hour, and then filtering the samples through a 2.5 μm Whatman grade 42 filter paper (Millipore Sigma, Burlington, MA, USA) to collect the extract. Concentrations of NH_4_^+^ and NO_3_^-^ (following reduction by cadmium) were determined based on absorption associated with indophenol blue (660 nm) and Griess azo dye (540 nm), respectively. PO_4_^3-^ test extracts were made using the Bray method by combining 3 g of sample with 25 mL of 0.1% NH_4_F and 0.2% HCl, shaking for 2-5 min, settling for an hour, and then filtering through a 2.5 μm (Whatman grade 42) to collect the extract. PO_4_^3-^ concentration was determined based on the absorption associated with molybdenum blue (660 nm). To measure pH, samples were prepared by combining 1 g ground sample and 1 mL deionized water.

### 2.4. DNA extraction and sequencing

We extracted nucleic acids from frozen bokashi samples using the Qiagen DNeasy PowerSoil kit (Qiagen, Germantown, MD, USA) and eluted in 60 μl of TBE buffer. Gel electrophoresis confirmed that sample extractions yielded amplifiable 16S rRNA gene and negative control (blank) extractions did not. DNA concentrations in all samples and negative controls were quantified using a NanoDrop ND-1000 (Thermo Scientific, Inc., Wilmington, DE, USA). The DNA extracts were sent to Integrated Microbiome Resource (IMR) at the Centre for Comparative Genomics and Evolutionary Bioinformatics (CGEB) at Dalhousie University for sequencing.

Bacterial and fungal diversity were assessed by sequencing the V4-V5 region of the 16S rRNA gene and the ITS region, respectively. All samples were prepared for sequencing following the Microbiome Amplicon Sequencing Workflow (Comeau et al., 2017). To sequence bacteria, the 515F 5’-GTGYCAGCMGCCGCGGTAA-3’ and 926R 5’-CCGYCAATTYMTTTRAGTTT-3 primers were used (Parada et al., 2016; Walters et al., 2015). To sequence fungi, the ITS86F 5’-GTGAATCATCGAATCTTTGAA-3’ and ITS4R 5’-TCCTCCGCTTATTGATATGC-3’ were used (Beeck et al., 2014). These primers had appropriate Illumina adapters and error-correcting barcodes unique to each sample to allow up to 380 samples to be simultaneously run per single flow cell. To reduce potential effects of PCR bias, each sample was amplified as an undiluted template and at a 1:10 dilution and pooled. All samples were normalized and PCR clean-up was performed using a high-throughput Charm Biotech Just-a-Plate 96-well Normalization Kit (Charm Biotech, Cape Girardeau, MO). The normalized samples were pooled together and the final library was quantified using a Qubit with PicoGreen (Invitrogen, United States). Finally, the library was sequenced on the Illumina MiSeq platform (Illumina, San Diego, CA) using 300+300 bp paired-end V3 chemistry, producing on average 55,000 raw reads per sample. Raw sequence files were deposited to the NCBI Sequence Read Archive under the BioProject accession number: PRJNA746996.

### 2.5. Processing and analyses of 16S rRNA gene and ITS sequence data

We processed sequences in QIIME 2 v.2020.8 (Bolyen et al., 2019), using a modified pipeline workflow developed by (Comeau et al., 2017). Primer sequences from both sequenced datasets were removed from sequenced data using the cutadapt plugin (Martin, 2011). Fastq reads were then filtered, trimmed and merged in DADA2 (Callahan et al., 2016) to generate a table of amplicon sequence variants (ASV). ASVs with a frequency of <0.1% of the mean sample depth were removed to account for a bleed-through between MiSeq runs, and the remaining reads were used for the subsequent analysis. For the 16S rRNA gene dataset, sequences belonging to mitochondria and chloroplasts were filtered out. For ITS data, one sample (Exp.1 - Control, Day 0) was excluded from downstream analyses as it only had 2,000 reads. A multiple-sequence alignment was created using MAFFT, and FastTree was used to create an unrooted phylogenetic tree, both with default values (Price et al., 2010). Taxonomy was assigned to the ASVs using a Naïve-Bayes classifier compared against a SILVA v 138.99 reference database trained on the 515-926 region of the 16S rRNA gene (Bokulich et al., 2018). Rarefaction curves showed that the majority of samples reached asymptote, indicating sequencing depth was appropriate for analyses in both bacterial and fungal datasets (Fig. S1).

### 2.6. Data analyses

We analyzed the effects of soil amendments on plant aboveground biomass and leaf chlorophyll content using linear models (separately for each plant species). In order to address our first question regarding the influence of bokashi on plant growth, we tested the specific contrast of a difference in means between the four experimental bokashi treatment levels and the soil and compost treatment levels. We used linear mixed models to examine univariate responses over the course of bokashi fermentation/development for nutrients, pH, and bacterial and fungal Shannon diversity. Independent fixed effects for these models were bokashi treatment type, time since the start of bokashi development, and their interaction. Individual experimental bokashi pile replicates were included as a random intercept effect to account for repeated measurements through time. We fit models separately for each of the two bokashi development experiments, as well as a third model to compare the controls across the two experiments. Models were fit using the lme4 package in R v 1.1-23 (Bates et al., 2015), and fixed effects tests were performed using the car package v 3.0-10 (Type II Wald chi-square tests).

To visualize the community composition, we constructed bubble plots and stacked bar plots of bacterial relative abundances using the ggplot2 package in R v 4.0.3. (Wickham et al., 2016). Alpha-diversity was estimated using Shannon diversity index (Shannon, 1948). Bray-Curtis dissimilarity was used to assess community structure (i.e., β-diversity). The clustering of bacterial communities was visualized via principal coordinate analysis (PCoA) plots constructed from Bray-Curtis distance matrices using ggplot2 (Wickham et al., 2016). We used Permutational Multivariate Analysis of Variance (PERMANOVA) tests to determine whether the bokashi type, time since the start of bokashi development, pH, or nutrients shaped microbial community structure (Anderson, 2001, 2017) using the adonis function from the R vegan package (Oksanen et al., 2016). We used Bray-Curtis dissimilarity with 1000 permutations. The strata term was used to limit permutations within individual experimental bokashi pile replicates to account for repeated measurements through time. We compared across both experiments, but we also ran PERMANOVAs separately for each of the two bokashi maturation experiments, as well as a third model to compare the Experiment 1 control to the Experiment 2 control.

## 3. Results

### 3.1. Growth experiment

Dry biomass and estimated chlorophyll concentration, for both kale and cucumber plants, were significantly higher for plants treated with bokashi than for those receiving compost or soil alone (Figure 2a,b; P < 0.001 for all contrast tests). The effect was more pronounced for kale, where mean biomass increased more than four fold when treated with bokashi relative to the soil-only or compost treatments. For both species, dry biomass was significantly related to variation in leaf chlorophyll associated with bokashi amendment (Figure 2b; kale *r^2^* = 0.51, cucumber *r^2^* = 0.54). The slope of this relationship was significantly greater for kale than for cucumber (0.017 vs. 0.006).

**Figure 2:**
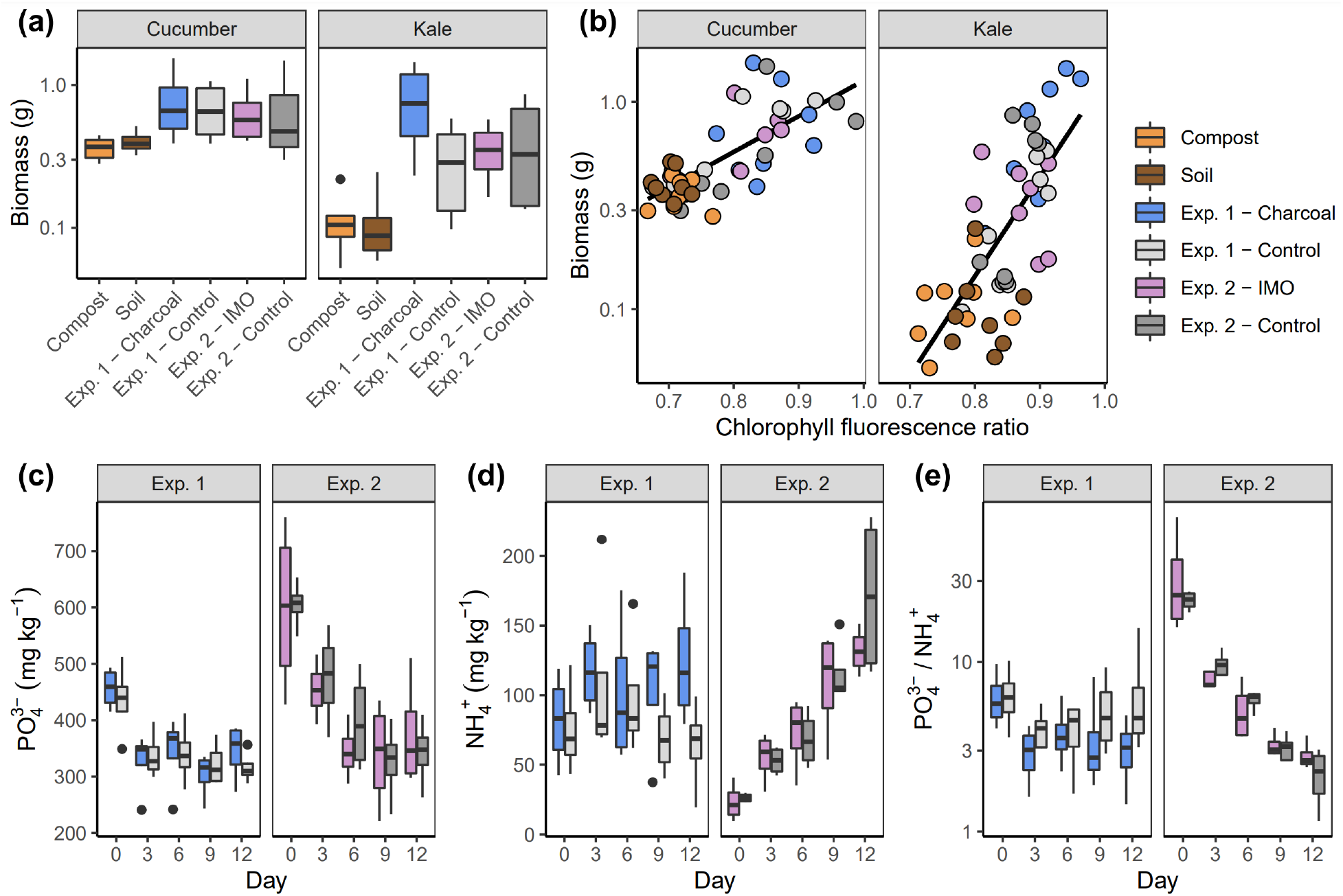
Bokashi nutrient composition and effect on plant growth. (a) Aboveground dry biomass of cucumber and kale plants treated with fully-matured (day 12) bokashi, compost, or soil. Plants grew for three weeks before collecting biomass. (b) Aboveground dry biomass and leaf chlorophyll fluorescence ratio (*F*_735_/*F*_700_) values of cucumber and kale plants treated with fully-matured bokashi, compost, or soil. (c) PO_4_^3-^ levels, (d) NH_4_^+^ levels, and (e) the ratio of PO_4_^3-^ levels to NH_4_^+^ levels in bokashi during the 12-day transformation period, shown separately for the two experiments.

### 3.2. Nutrient analysis

PO_4_^3-^ concentrations across all bokashi types exhibited a decrease over time for all bokashi types (Figure 2c). The PO_4_^3-^ concentration for bokashi made in Exp. 2—both IMO and Control—was initially higher (between days 0 to 3) than those made in Exp. 1 (median starting values of 608 mg/kg vs. 440 mg/kg). This aligns with the PO_4_^3-^ concentrations of the manure used: 1810 mg/kg for Exp. 2 vs. 680 mg/kg for Exp. 1. On the final day of bokashi maturation (Day 12) there was no significant difference between the PO_4_^3-^ concentrations of the different bokashi types (P = 0.645).

NH_4_^+^ concentrations exhibited variable trends over time by type (Figure 2d). NH_4_^+^ levels in bokashi made in Exp. 2—both IMO and Control—increased consistently over time, whereas NH_4_^+^ levels in Exp. 1 – Charcoal mostly increased but dipped slightly on Day 6 before rising again, and levels in Exp. 1 – Control peaked on Day 3 before decreasing steadily. Though both experimental controls contained the same ingredients—but varied in time during which they were made and, thus, when the manure was collected—they exhibited different trends in NH_4_^+^ levels over time and variable final NH_4_^+^ amounts (P = 0.004). The NH_4_^+^ concentration of the manure used in Exp.1 was 90.4 mg/kg and for the manure in Exp. 2 was 133 mg/kg. NO_3_^-^ concentrations remained constant over time across all bokashi types and were generally negligible (< 3 mg/kg) (Figure S3). A comparison of the trends in NO_3_^-^ and NH_4_^+^ levels over time illustrates no relationship between the concentrations of the two inorganic forms of N in any of the bokashi types.

Overall, the two different experiments exhibited distinct patterns of nutrient dynamics, with far more variation through time in Exp. 2 as PO_4_^3-^ decreased and NH_4_^+^ increased. Regardless of treatments, Exp. 2 began with substantially higher ratios of PO_4_^3-^ to NH_4_^+^ (median of 23.5 vs. 6.1 in Exp. 1), though both experiments converged on relatively similar ratios after 12 days of development (median of 2.6 for Exp. 2 vs. 3.8 for Exp. 1). (See Table A.1 for statistical tests for all nutrients).

### 3.3. Microbial analysis

To examine the relationships between bacterial and fungal communities and the bokashi type, bokashi maturation time, and nutrient profiles, we sequenced the 16S rRNA genes (for days 0, 3, 6, 9 and 12) and ITS regions (for days 0, 6, and 12) from the bokashi metagenomic DNA (Figure 1, Table S2, Table S3). Rarefaction curves generated from all samples generally approached saturation suggesting that both the fungal and bacterial communities were adequately sampled (Figure S1).

We used Bray-Curtis beta diversity as a metric to determine the effects of different bokashi treatments and maturation on shaping overall community structure for both bacterial and fungal communities. Principal coordinate analysis based on Bray-Curtis distances between amplicon sequence variants (ASVs) showed separation by Exp. 1 and 2 and days of maturation (Figure 3 a,b). For the 16S rRNA gene amplicon data, bacterial communities for all experimental types clustered on Day 0 (Figure 3a, lower right quadrant), and then separated into two groups by clustered by experiment: Exp. 1 - Charcoal and Exp. 1 - Control data points clustered together and away from the Exp. 2 - IMO and Exp. 2 - Control cluster (Figure 3a). The fungal communities were more similar between the two experiments and their community changed over time (Figure 3b).

**Figure 3:**
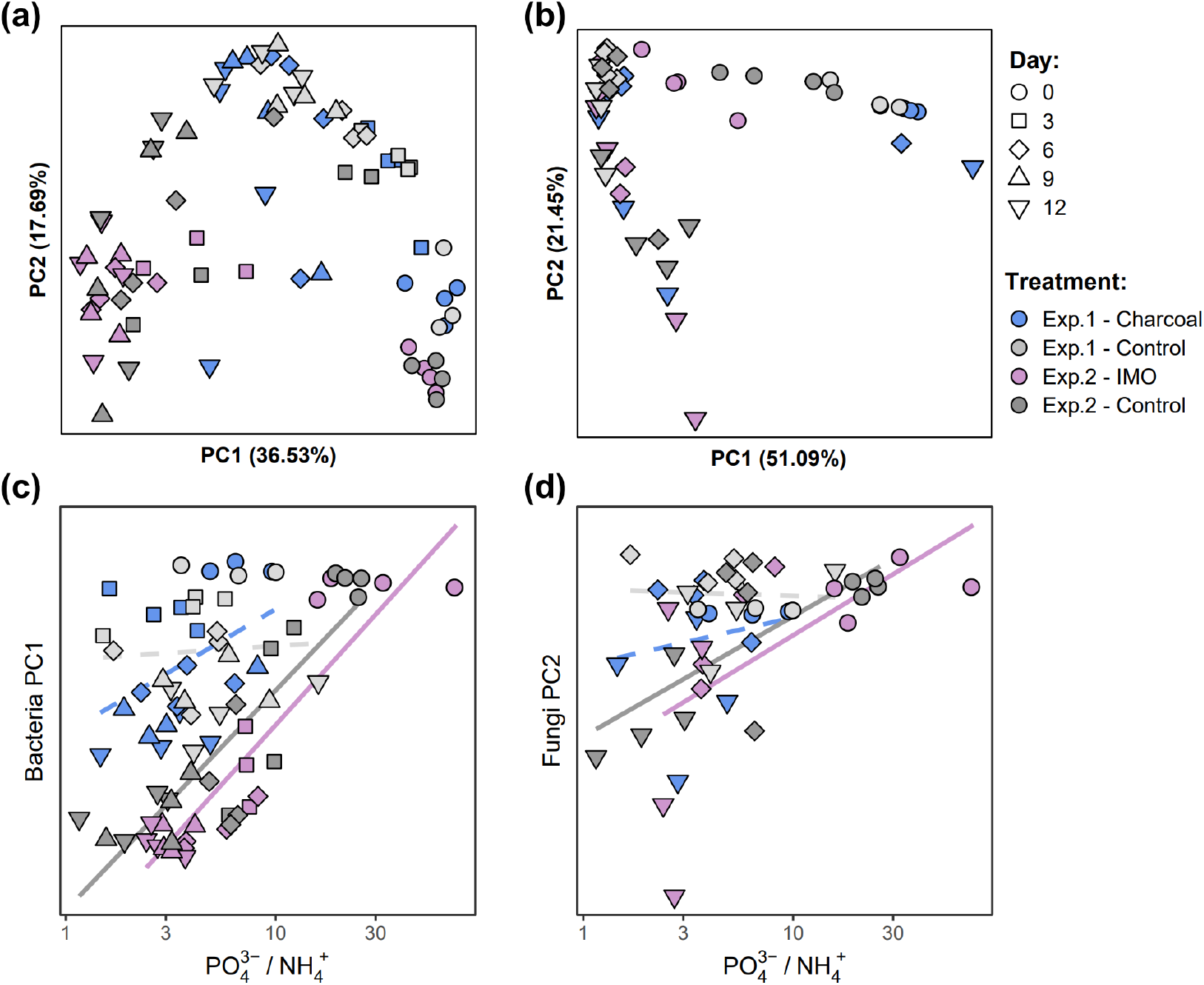
Principle coordinate analysis (PCoA) of the microbial community composition throughout bokashi maturation based on Bray-Curtis distances. (a) Bacterial microbial community structure (16S rRNA gene amplicon data) and (b) fungal microbial community structure (ITS amplicon data). Samples are color coded based on the experimental treatment and shapes indicate days of maturation when samples were collected. Ratio of PO_4_^3-^/NH_4_^+^ values plotted against the (c) Bacteria PCoA Axis 1 (from panel (a)), and (d) Fungi PCoA Axis 2 (from panel(b)). Samples are color coded based on the experimental treatment. Solid lines indicate slopes that are significantly different from zero (P < 0.05), and dashed lines indicate nonsignificant slopes (P > 0.05).

To assess variance partitioning between different bokashi treatments and maturation, we ran PERMANOVAs separately for each of the two bokashi experiments. PERMANOVA analyses with experimental treatment and time for each experiment and controlled for repeated measurements through time indicated significant differences (p < 0.001) for both treatment and time (Table S4). There were no significant differences for interaction terms between time and treatment. Both bacterial and fungal communities were more strongly partitioned by time than treatment. Time alone accounted for 28.1% and 36.5% for bacterial communities in Exp. 1 and Exp. 2, respectively (Table S4). For the fungal communities, time accounted for 34.4% and 31.4% for Exp. 1 and Exp. 2, respectively. The different treatments for each experiment explained only between 4.1 and 7.5% of the variation in community structure across samples (Figure 3 a, b, Table S4).

To explore the link between nutrients and microbial communities, we plotted the values of the PO_4_^3-^ to NH_4_^+^ ratio against the principal coordinate axis (Figure 3 c, d). The two experiments, and thus the two controls, differed in one ingredient - the manure which contained different starting amounts of NH_4_^+^. The change in microbial community composition for both fungal and bacterial communities in Exp. 2 was associated with the changes in the ratio of PO_4_^3-^ and NH_4_^+^ (and bokashi maturation), but this was not the case for Exp. 1 (Figure 3 c, d).

We next examined the changes in bacterial community composition by 16S rRNA gene amplicon sequencing across different bokashi treatments over time and investigated whether the source of microorganisms in the bokashi was from the soil or manure.

Bacterial communities across all bokashi types were similar, the four most dominant phyla included Proteobacteria (66% relative average abundance), Bacteroidota (21%), Firmicutes (7%) and Acidobacteriota (2.8%) (Table S2, Figure S6). At the phylum and order level of classification, all experiments had similar percentages of different taxonomic groups. However, at the genus level, Experiment 1 differed significantly from Experiment 2. Most notably, relative abundances of the genus *Thermomonas* (phylum Proteobacteria, family Xanthomonadaceae) were higher across Experiment 1, while *SN8* (a different genus within the Xanthomonadaceae family) and *Taibaiella* (phylum Bacteroidota, family Chitinophagaceae) were higher in Experiment 2.

We observed an increase of 8.3%, 4.8%, and 3.7% in average relative abundances of *Dyella, Luteibacter*, and *Edaphobacter*, respectively, in all bokashi types over time (Figure 4a, Figure S6). These bacteria began at low relative abundances in the soil. Other trends across all bokashi types included increases in relative abundances of *Allorhizobium/Rhizobium, Azotobacter, Burkholderia/Paraburkholderia, Chitinophaga, Gluconacetobacter, Paenibacillus, Pseudoxanthomonas, Taibaiella* (more so in Exp. 2 bokashi), and *Thermomonas* (more so in Exp. 1), and slight increases in *Anaerosinus, Myxococcaceae, Rhodanobacter, Rhondanobacteraceae*, and *Sphingomonas. Clostridium sensu stricto* and *Dokdonella* relative abundances increased in Exp. 1 bokashi, but remained stable in Exp. 2. *Dokdonella* was present in the soil, suggesting it may have been the initial population. *Stenotrophomonas* relative abundance was very high in the IMO ingredient and high in the manure used in the Exp. 1; in all bokashi types, *Stenotrophomonas* spiked by Day 3 before decreasing until the end of maturation. *Xanthomonas massiliensis*, strain SN8, only appeared in bokashi sampling on the final day of maturation in Exp. 1, but increased substantially during maturation in Exp. 2 bokashi. SN8 was present in the manure used in Exp. 2 (and not in the manure of Exp. 1) and in the IMO. Relative abundances of taxa belonging to *Pseudomonas, Flavobacterium, Comamonas, Comamonadaceae, Brevundimonas*, and *Acinetobacter* decreased in all bokashi types during maturation. All of these bacteria were present in both manures and the IMOs, but only *Comamonadaceae* was also in the soil.

**Figure 4:**
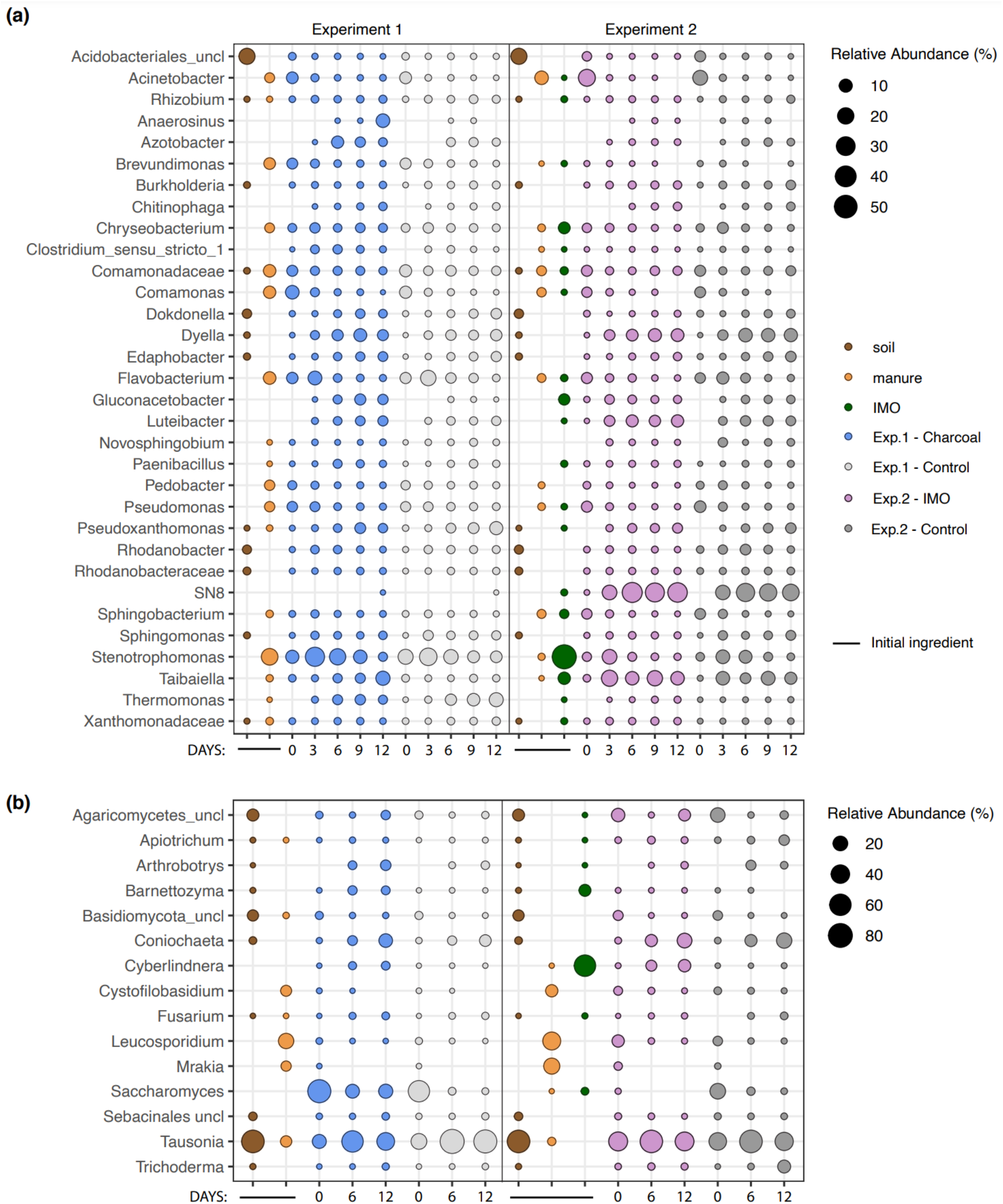
Bubble plots of the relative abundance (%) of the genus-level taxa during bokashi maturation compared between two experiments. (a) Relative abundances of bacteria (16S rRNA gene amplicon) in each bokashi type during days 0, 3, 6, 9, and 12 of maturation. (b) Relative abundances of fungi (ITS amplicon) in each bokashi type during days 0, 6, and 12 of maturation. Soil was identical for both experiments; manure was sourced at different times. Circle area represents relative abundances of a taxon and absence of a circle indicates a relative abundance of <0.01%. Only taxa with average relative abundance of >0.5% across all treatments are shown.

Analysis of fungal populations by ITS sequencing revealed high relative abundance of Basidiomycota fungi in the initial soil and manure that decreased slightly during bokashi maturation. Ascomycota fungi started with a lower relative abundance in starting soil and manure, but increased during the bokashi maturation process. The Basidiomycota fungi in the Sebacinales order and *Tausonia* genus were present in the initial soil and manure, and both maintained their relative abundance throughout the bokashi maturation process (Figure 4b). *Apiotrichum* was stable in abundance in Exp. 1 - Charcoal bokashi and increased in overall average relative abundance in Exp. 2 - IMO. *Cystofilobasidium, Leucosporidium, and Mrakia* were introduced at high levels from the manure, but the relative abundance of each decreased during bokashi maturation. Among Ascomycota, the relative abundance of *Arthrobotrys* and fungi in the class Sodariomycetes, including *Coniochaeta, Fusarium*, and *Trichoderma*, gradually increased in all bokashi types (Figure 4b). The relative abundance of Saccharomycetes genera *Barnettozyma* and *Cyberlindnera* increased in both the Exp. 1 - Charcoal and Exp. 2 - IMO experimental bokashi groups, but remained stable in control bokashi. *Saccharomyces* abundance decreased during the bokashi maturation period for all experimental and control groups.

Alpha diversity, measured as both species richness (the number of different taxa in a sample) and evenness (the distribution of abundances of different taxa), was higher on Day 0 in Experiment 2 for both bacterial and fungal communities. While the Shannon diversity in Exp. 1 bokashi types increased from Day 0 to Day 12, in Exp. 2 bokashi types, there appeared to be an initially high Shannon richness on Day 0, followed by a decrease by Day 6, and finally an increase resembling a return to Day 0 levels on Day 12 (Figure 5 a,b). Shannon diversity correlated with the ratio of PO_4_^3-^ and NH_4_^+^ for bacteria only in Experiment 2. There was no correlation between the Shannon diversity index calculated for fungi for any bokashi treatments.

**Figure 5:**
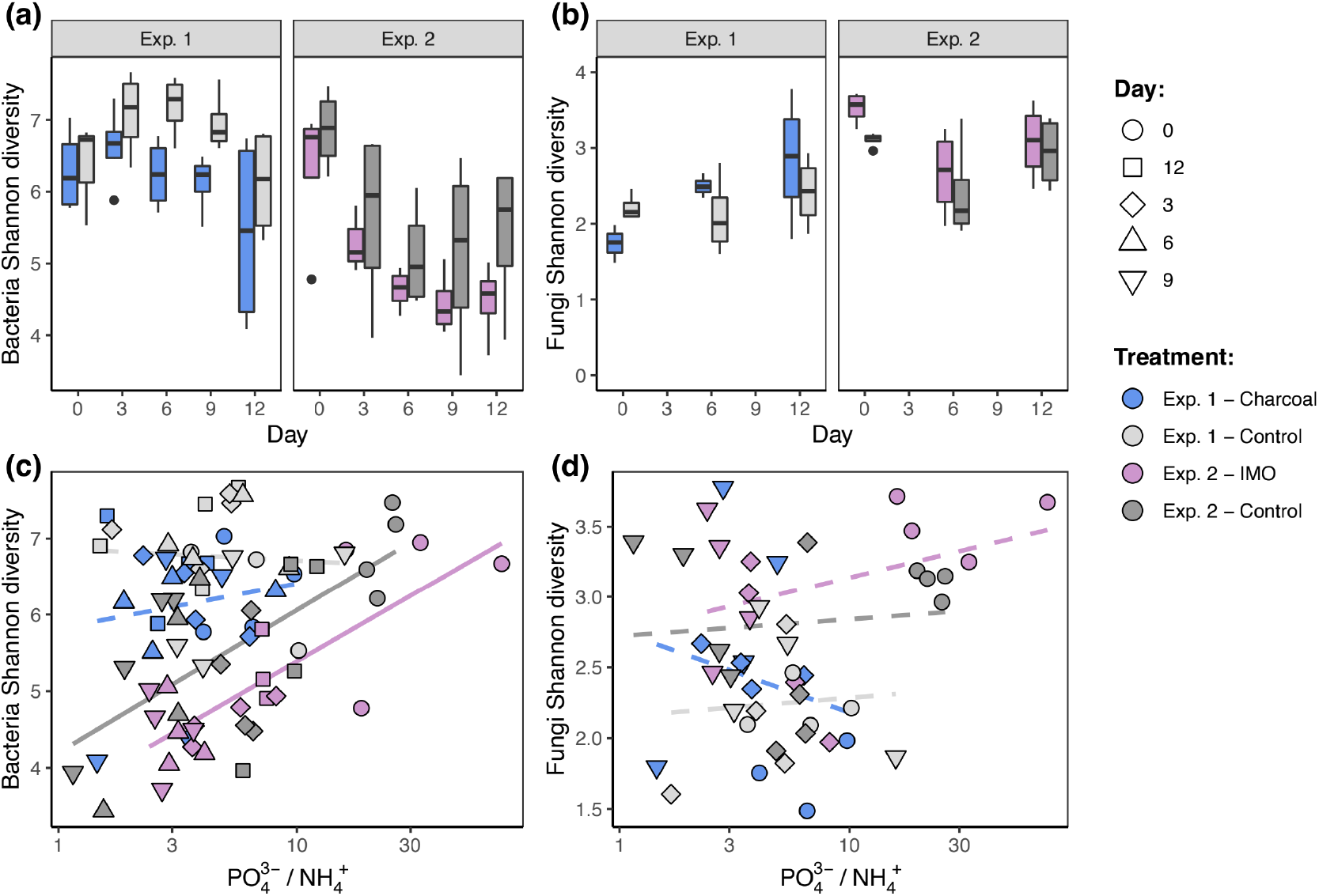
Alpha diversity of bokashi microbial communities during maturation. Shannon richness of (a) bacterial communities during days 0, 3, 6, 9, and 12 of bokashi maturation and (b) fungal communities during days 0, 6, and 12 of bokashi maturation between two experiments. Shannon richness by PO_4_^3-^ to NH_4_^+^ ratio for (c) bacterial communities and (d) fungal communities. Solid lines indicate slopes that are significantly different from zero (P < 0.05), and dashed lines indicate nonsignificant slopes (P > 0.05).

## 4. Discussion

Increasing agricultural pressures on the environment raise the urgency of using agroecological approaches that promote dynamic nutrient cycling, minimize the negative impact of high conventional fertilizer inputs, and meet yield demands. Bokashi, a readily available, easy-to-prepare organic amendment, can improve soil composition by increasing concentrations of bioavailable phosphorus and nitrogen pools while also contributing beneficial microbes. More environmentally and economically sustainable agricultural techniques can be promoted by understanding the microbial and nutrient inputs of agroecological techniques, like the traditional Japanese fertilizer bokashi. Here, we report on the efficacy of bokashi made from variable starting materials and on the nutrient and microbial transformation of the fertilizer during maturation.

Cucumber and kale seedlings grown in bokashi-amended soils grew significantly larger and had higher leaf chlorophyll concentrations compared to seedlings grown in compost-amended soils or in soil alone (Figure 2a). Our finding that bokashi improves plant growth is consistent with previous studies (Álvarez-Solís et al., 2016; Aurora Gomez-Velasco et al., 2014; Bautista-Cruz et al., 2014; Boechat et al., 2013; França et al., 2016; Jaramillo-López et al., 2015; Lima et al., 2015; Peralta-Antonio et al., 2014). The positive correlation between bokashi amendments, leaf chlorophyll concentration, and plant biomass further suggests a pathway for enhanced plant growth by bokashi through increased bioavailable N concentrations (Figure 2b), as does the positive correlation between NH_4_^+^ concentrations of bokashi and chlorophyll concentrations in plant leaves (Fig. S5). Trends in NH_4_^+^ and PO_4_^3-^ concentrations differed between experiments, which may have been due to the fact that because the experiments were conducted at different times, and the manures used were collected and stored at different times. Regardless, the final ratios of PO_4_^3-^ to NH_4_^+^ were broadly similar across all four bokashi treatments. This may suggest some degree of deterministic processes in microbial function and succession, as community composition was observed to correlate with PO_4_^3-^ / NH_4_^+^ (particularly so in Exp. 2 where PO_4_^3-^ / NH_4_^+^ exhibited a significant decrease throughout maturation; Figure 3).

Analysis of the bacterial populations reveals increases in abundances of plant growth promoting rhizobacteria (PGPR) genera, many of which have been linked to improved plant growth (Babalola, 2010; Grobelak et al., 2015; Kaymak et al., 2009). Of the PGPR, we identified increasing abundances of nitrogen-fixers such as *Allorhizobium/Rhizobium, Azotobacter, Burkholderia/Paraburkholderia*, and *Paenibacillus* (Estrada-De Los Santos et al., 2001; Govindarajan et al., 2008; Martín et al., 1993; von der Weid et al., 2002; Wang et al., 2020; Zahran, 1999), which could have contributed to increasing concentrations of NH_4_^+^ in mature bokashi. The relative abundance of several other nitrogen-fixing bacterial genera also increased, including *Clostridium sensu stricto, Dokdonella, Rhodanobacter*, and *Sphingomonas* (Figueroa-González et al., 2016; Huang et al., 2016; Wang et al., 2020; Zhou et al., 2021). Conversely, phosphate solubilizing bacteria (PBR) such as *Acinetobacter* and *Pseudomonas* decreased in abundance, which was consistent with decreased concentrations of PO_4_^3-^ during bokashi maturation (Pathma and Sakthivel, 2013). Such microbially-mediated changes likely improve the efficacy of bokashi by reducing nutrient forms that are more readily leached from soils, such as the anions NO_3_- and PO_4_^3^-. Once applied as a soil amendment, nitrogen and phosphorus can become more readily available for plant uptake via nitrification and mineralization, respectively.

We also observed increasing relative abundances of bacterial taxa associated with environmental decomposition of plant biomass, including *Chitonophaga, Dyella, Novosphingobium, Pseudoxanthomonas, Taibaiella*, and *Thermomonas* (Akyol et al., 2019; Cai et al., 2018; Cecil et al., 2018; Larsbrink et al., 2017; Liang et al., 2014; Zainudin et al., 2013). Their observed increase in abundance during bokashi maturation could be due to the cellulose, hemicellulose, and lignin-rich composition of rice hulls used in bokashi preparation (Stroeven et al, 1999).

Similarly, decomposing fungi from both Ascomycota and Basidiomycota divisions increased in relative abundance during bokashi maturation. The soil used to make all bokashi types had a high abundance of *Tausonia*, accounting for the enhanced representation of these fungi in all of the bokashi types on Day 0. *Tausonia* is commonly found in decaying leaf litter and produces cellulase, and its relative abundance increased during the bokashi maturation process (Dennis, 1972; Jimenez et al, 1991; Mestres et al, 2011). Members of the class Sodariomycetes, including *Coniochaeta, Fusarium*, and *Trichoderma*, contribute to breakdown of cellulose and lignin in soil isolates and leaf litter (Bhatnagar et al., 2018; Jäger et al., 2011; U’Ren and Arnold, 2016). Additionally, members of the order Saccharomycetes such as *Cyberlindnera* and *Barnettozyma* produce xylanase and cellulase, respectively, and both genera have been associated with degrading leaf litter and wood (Guamán-Burneo et al., 2015; Morais et al., 2013). Interestingly, another genus within this order, *Saccharomyces*, that does not produce these enzymes had a decrease in relative abundance during bokashi maturation, despite being added on Day 0 to both experimental controls and Exp. 1 - Charcoal. This suggests that *Saccharomyces* was outcompeted by other taxa during the maturation process.

Increases in the relative abundance of other fungal taxa could contribute to previously described pathogenic pest control benefits to bokashi (Lwin and Ranamukhaarachchi, 2006; Olle, 2020). In addition to degrading cellulose, *Arthrobotrys* species have been linked to trapping plant parasitic soil nematodes, which in turn promotes improved root vasculature (Lan et al., 2016, p. 20; Niu and Zhang, 2011).

Additionally, members of the bacterial genus *Paenibacillus* have demonstrated antagonistic activity toward phytopathogenic bacteria and fungi through generation of antimicrobial substances (von der Weid et al, 2003).

Many of the beneficial bacteria we observed in bokashi were initially present in the soil, suggesting the importance of the quality of the starting ingredients. Despite variation in bokashi microbial communities, all types of bokashi performed similarly in improving plant growth, thus, even as starting ingredients varied, we found that bokashi is an effective and adaptable fertilizer. The generation of ammonium rather than the more labile nitrate, the biological immobilization of phosphorus, the abundance of PGPR, and the potential pathogenic-controlling properties of bokashi suggest several advantages of this fertilizer over agrochemical fertilizers: the biochemical composition of bokashi may reduce nutrient runoff and foster beneficial microbial communities when used in agroecosystems. Future research should elucidate the biological pathways during bokashi maturation that lead to the observed nutrient concentrations; a study of the metagenomes and finer level characterization of isolates may shed light on pathogenic and beneficial microbes in promoting plant growth.

## Acknowledgements

This research was funded by the Pamela Daniels Fellowship and the Environmental Studies Department at Wellesley College (both awarded to NAS). We thank Sophie Rowland for helping with sample processing, the Asian Rural Institute (Tochigi, Japan) for teaching NAS about bokashi, and the Natick Community Organic Farm for providing manure.

