## Supplementary materials for "Microbial transformation of traditional fermented fertilizer bokashi alters chemical composition and improves plant growth"

**Figure S1.** Rarefaction curves for all samples for (a) 16S rRNA gene and (b) ITS region datasets.

**Figure S2.** Visual comparison of (a and b) cucumber and (c and d) kale growth after three weeks in each treatment. Plants on the left in each photo were grown in soils amended with bokashi (Exp. 1 - Charcoal); plants in the center in soil amended with compost; and plants on the right were grown in soil alone. Photos (a) and (c) were taken from the side and (b) and (d) were taken from the top of the same plants.

**Figure S3.** $NO_3^-$ levels in bokashi during the 12-day transformation period, shown separately for the two experiments.

**Figure S4.** pH of bokashi during the 12-day transformation period, shown separately for the two experiments.

**Figure S5.** $NH_4^+$ concentration in bokashi used to amend soil vs. (a) the chlorophyll fluorescence ratio of leaves and (b) aboveground biomass of cucumber and kale plants after three weeks of growth. Shading indicates the 95% confidence interval for slope parameters.

**Figure S6: Reproducible relative abundances and variation of bacterial communities during bokashi maturation as compared across two experimental conditions.** Stacked bar plots showing the relative frequency of 16S rRNA gene sequences assigned to each bacterial (a) phylum, (b) order, and (c) genus across samples. Samples are grouped by experiments and then by days (0, 3, 6, 9 and 12) of maturation. Each color represents a phylum in (a), order in (b)  or genus in (c). Only taxa >1% are shown; all others are grouped into Other. Replicates for each day/experimental condition are grouped within a panel.

**Figure S7: Reproducible relative abundances and variation of fungal communities during bokashi maturation as compared across two experimental conditions.** Stacked bar plots showing the relative frequency of ITS region sequences assigned to each fungal (a) phylum, (b) order, and (c) genus across samples. Samples are grouped by experiments and then by days (0, 6, and 12) of maturation. Each color represents a fungal phylum in (a), order in (b)  or genus in (c). Only taxa >1% are shown; all others are grouped into Other. Replicates for each day/experimental condition are grouped within a panel.

**Table S1.** P-values obtained by running linear mixed models to examine univariate responses over the course of bokashi fermentation/development for nutrients, pH, and bacterial and fungal Shannon diversity. Independent fixed effects for these models were bokashi treatment type, time since the start of bokashi development, and their interaction. Replicate bokashi piles were included as a random effect and the estimate standard deviation is reported (all response variables were standardized in units of SD).

**Table S2.** Relative ASV abundances (%) for bacterial data for each sample used in the bokashi study. Associated metadata for each sample are provided.

**Table S3.** Relative ASV abundances (%) for fungal data for each sample used in the bokashi study.  Associated metadata for each sample are provided.

**Table S4.** PERMANOVA results of 16S and ITS datasets from bokashi maturation experiments using the Bray-Curtis dissimilarity matrix. Variance explained by each factor.

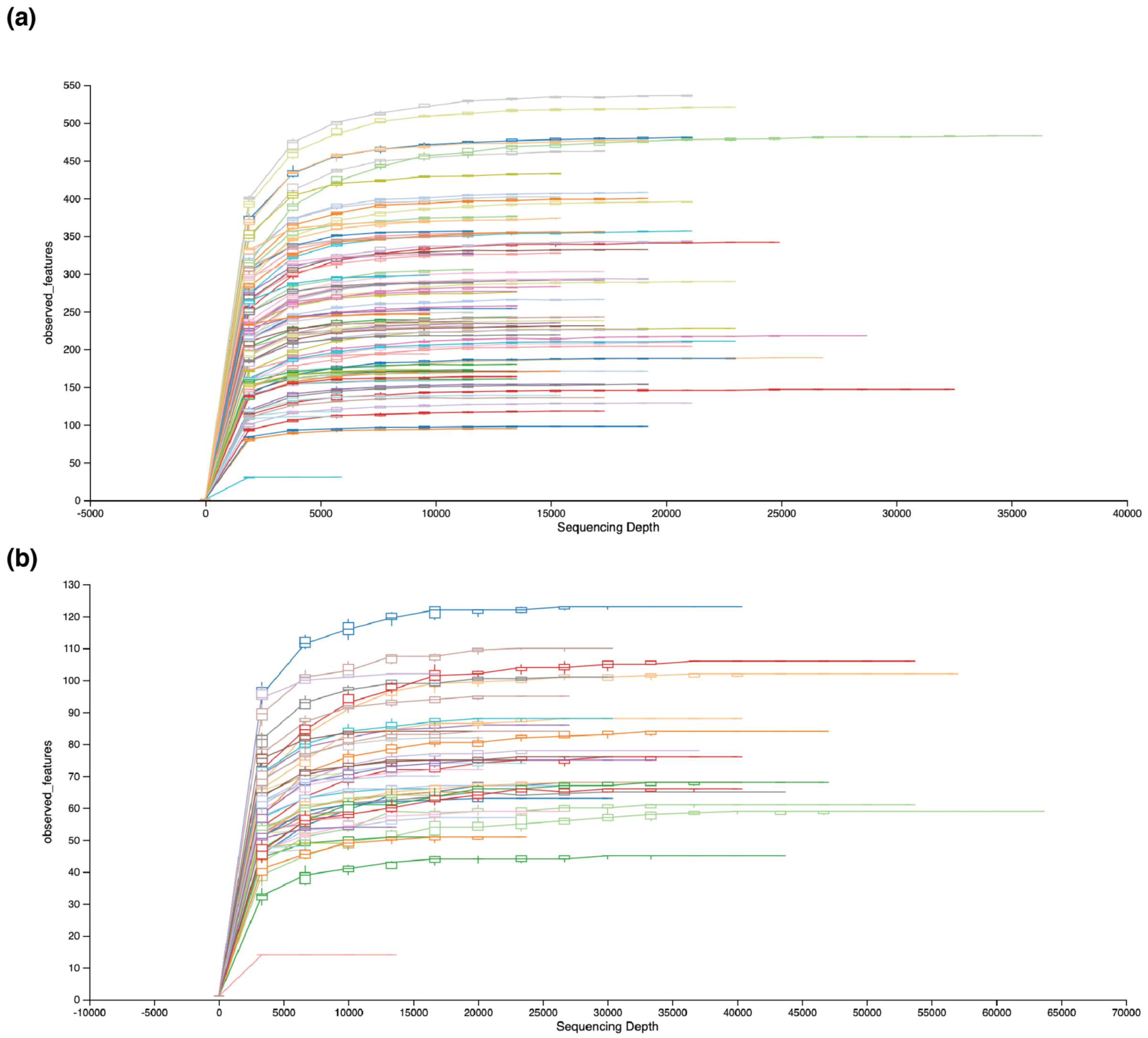


**Figure S1.** Rarefaction curves for all samples for (a) 16S rRNA gene and (b) ITS region datasets.

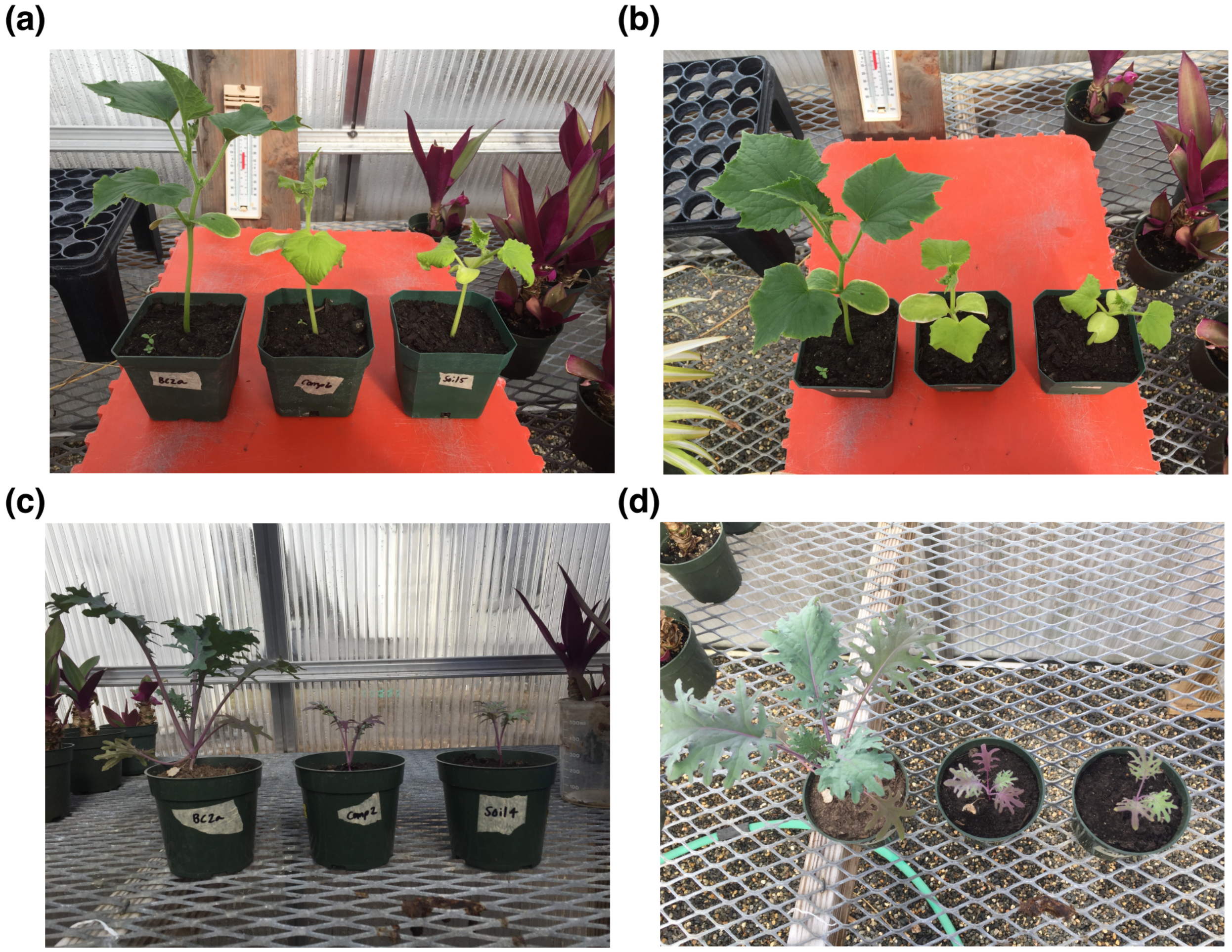


**Figure S2.** Visual comparison of (a and b) cucumber and (c and d) kale growth after three weeks in each treatment. Plants on the left in each photo were grown in soils amended with bokashi (Exp. 1 - Charcoal); plants in the center in soil amended with compost; and plants on the right were grown in soil alone. Photos (a) and (c) were taken from the side and (b) and (d) were taken from the top of the same plants.

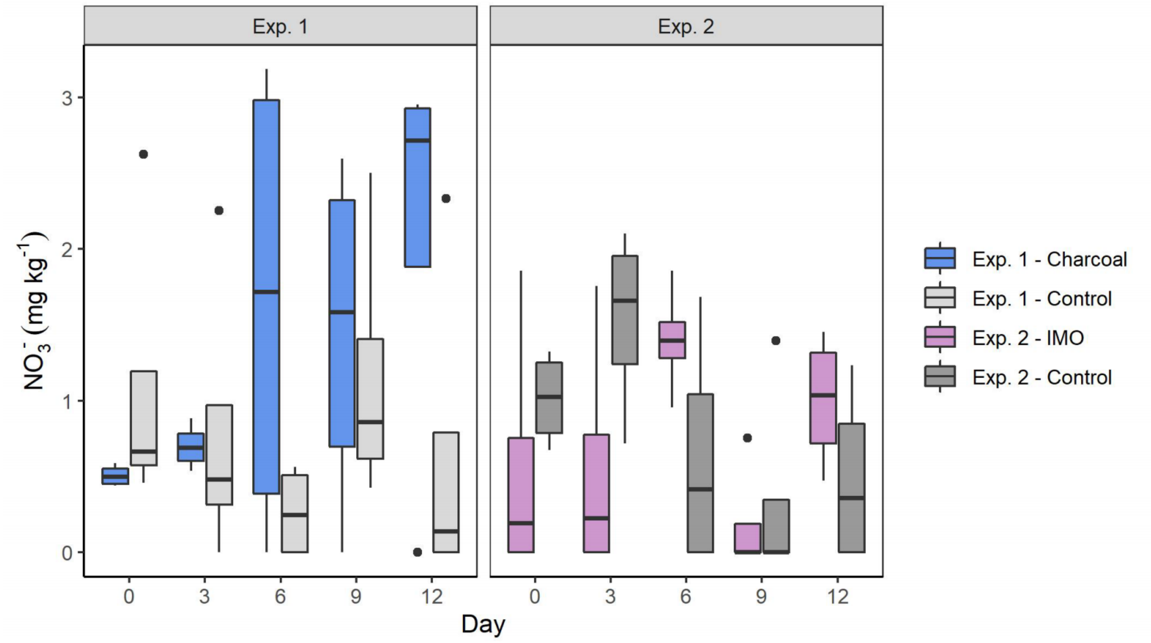


**Figure S3.** $NO_3^-$ levels in bokashi during the 12-day transformation period, shown separately for the two experiments.

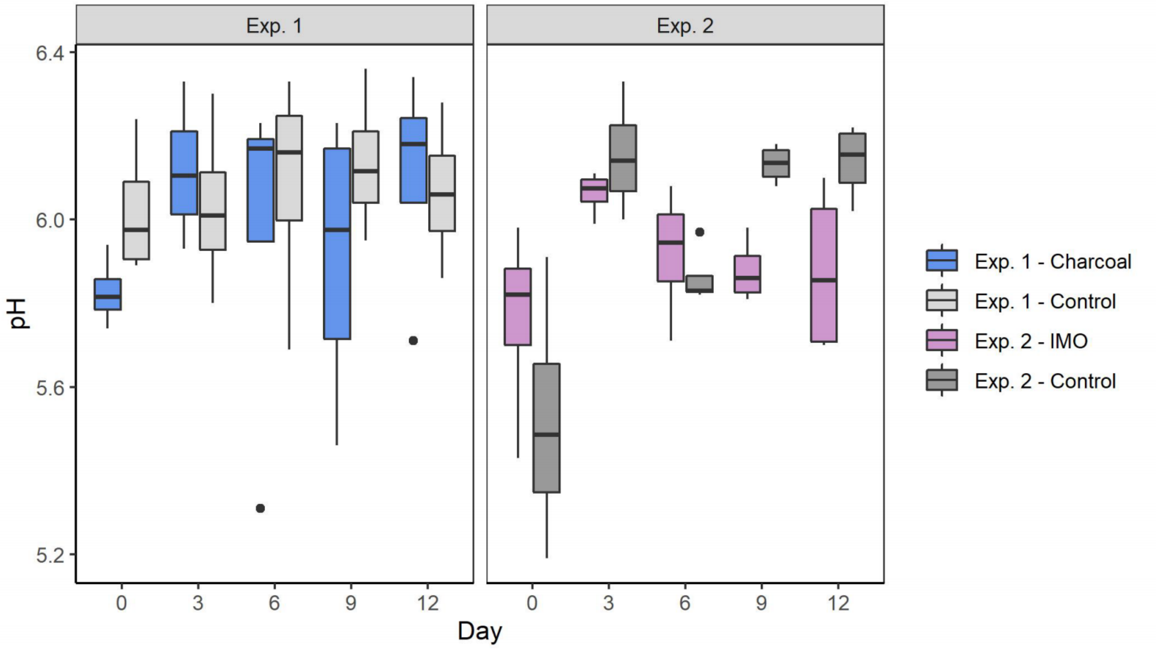


**Figure S4.** pH of bokashi during the 12-day transformation period, shown separately for the two experiments.

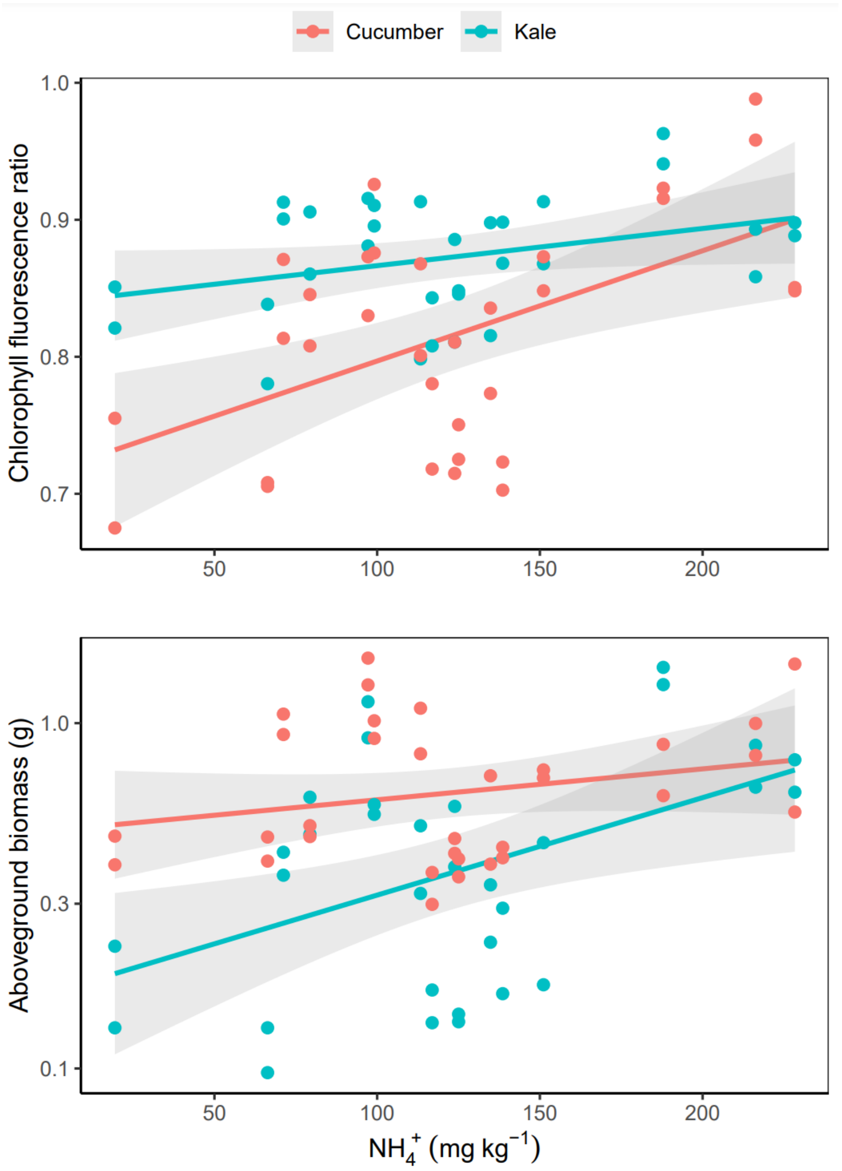


**Figure S5.** $NH_4^+$ concentration in bokashi used to amend soil vs. (a) the chlorophyll fluorescence ratio of leaves and (b) aboveground biomass of cucumber and kale plants after three weeks of growth. Shading indicates the 95% confidence interval for slope parameters.

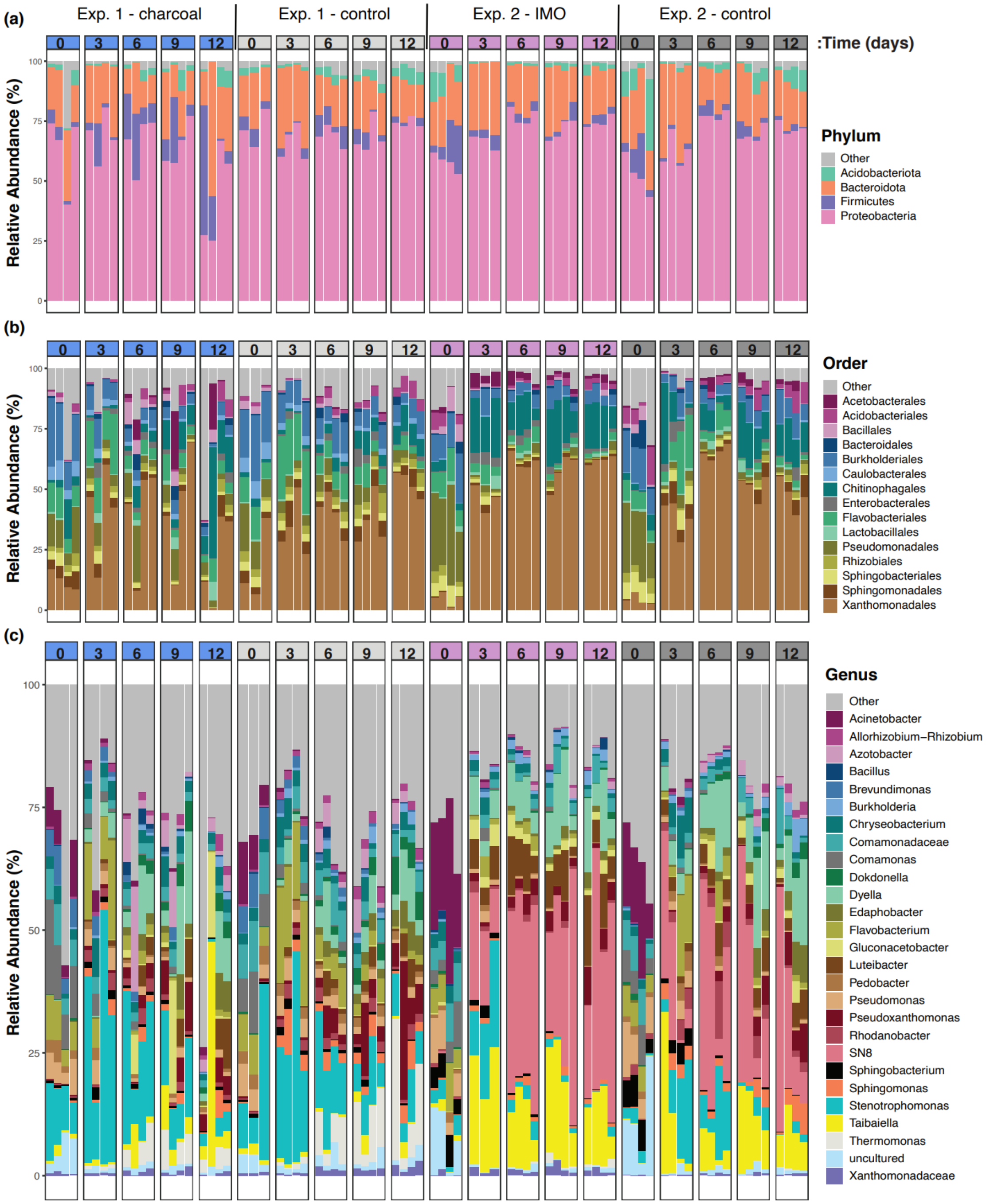


**Figure S6: Reproducible relative abundances and variation of bacterial communities during bokashi maturation as compared across two experimental conditions.** Stacked bar plots showing the relative frequency of 16S rRNA gene sequences assigned to each bacterial (a) phylum, (b) order, and (c) genus across samples. Samples are grouped by experiments and then by days (0, 3, 6, 9 and 12) of maturation. Each color represents a phylum in (a), order in (b) or genus in (c). Only taxa >1% are shown; all others are grouped into Other. Replicates for each day/experimental condition are grouped within a panel.

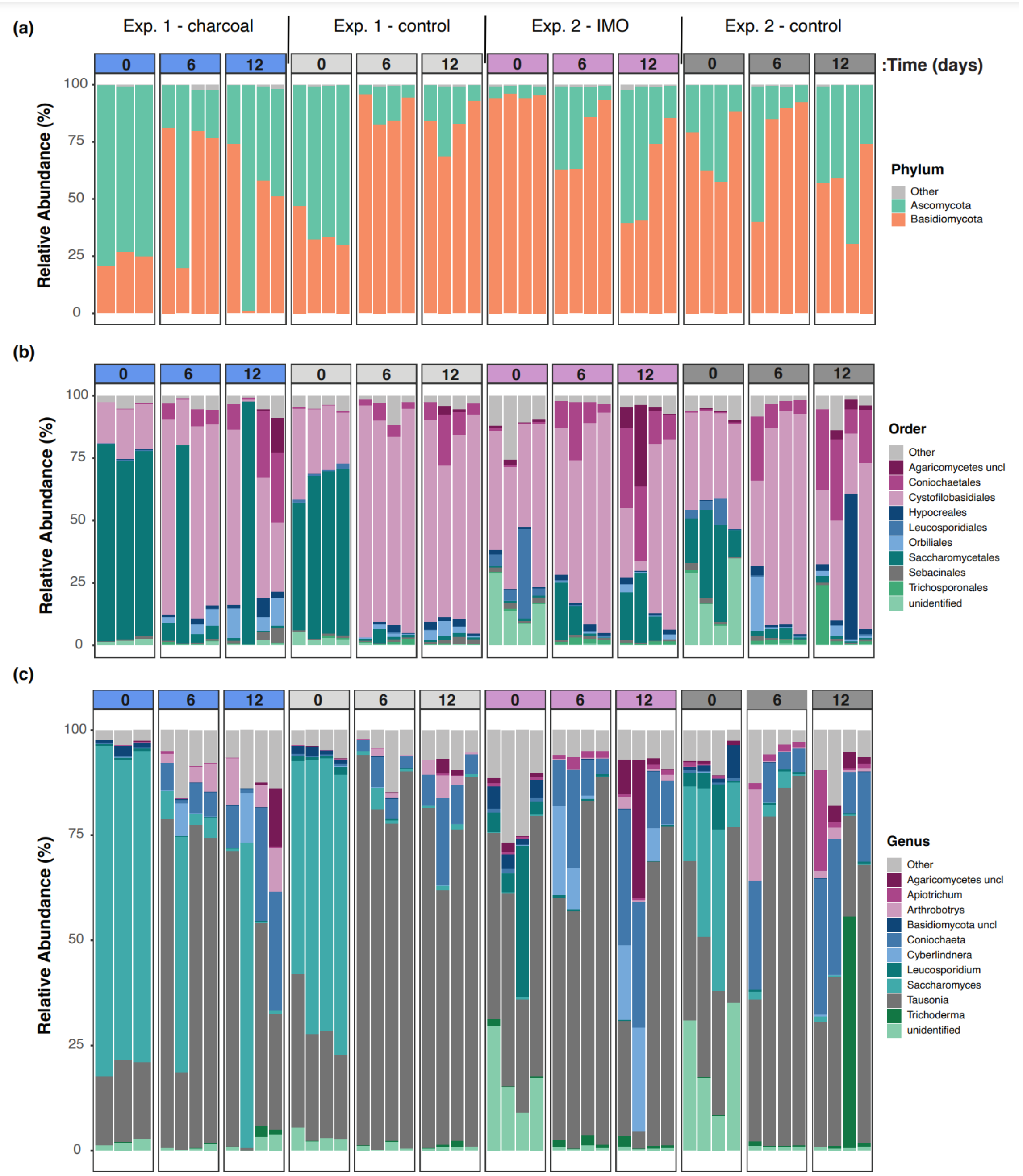


**Figure S7: Reproducible relative abundances and variation of fungal communities during bokashi maturation as compared across two experimental conditions.** Stacked bar plots showing the relative frequency of ITS region sequences assigned to each fungal (a) phylum, (b) order, and (c) genus across samples. Samples are grouped by experiments and then by days (0, 6, and 12) of maturation. Each color represents a fungal phylum in (a), order in (b) or genus in (c). Only taxa >1% are shown; all others are grouped into Other. Replicates for each day/experimental condition are grouped within a panel.

**Table S1.** P-values obtained by running linear mixed models to examine univariate responses over the course of bokashi fermentation/development for nutrients, pH, and bacterial and fungal Shannon diversity. Independent fixed effects for these models were bokashi treatment type, time since the start of bokashi development, and their interaction. Replicate bokashi piles were included as a random effect and the estimate standard deviation is reported (all response variables were standardized in units of SD).

| Model* | Treatment | Time | Treatment × Time | Replicate VC (SD) |
|---|---|---|---|---|
| ***$PO_4^{3-}$*** | | | | |
| Exp. 1 - Charcoal vs. Control | 0.874 | **< 0.001*** | 0.963 | 0.21 (0.46) |
| Exp. 2 - IMOs vs. Control | 0.881 | **< 0.001*** | 0.504 | 0.22 (0.47) |
| Exp. 1 Control vs. Exp. 2 Control | **0.025*** | **< 0.001*** | **< 0.001*** | 0.18 (0.43) |
| ***$NH_4^+$*** | | | | |
| Exp. 1 - Charcoal vs. Control | 0.170 | 0.935 | 0.132 | 0.11 (0.34) |
| Exp. 2 - IMOs vs. Control | 0.363 | **< 0.001*** | 0.149 | 0.02 (0.14) |
| Exp. 1 Control vs. Exp. 2 Control | 0.816 | **< 0.001*** | **< 0.001*** | 0.10 (0.31) |
| ***$PO_4^{3-}/NH_4^+$*** | | | | |
| Exp. 1 - Charcoal vs. Control | 0.250 | 0.316 | 0.155 | 0.20 (0.45) |
| Exp. 2 - IMOs vs. Control | 0.539 | **< 0.001*** | 0.692 | 0.02 (0.15) |
| Exp. 1 Control vs. Exp. 2 Control | 0.352 | **< 0.001*** | **< 0.001*** | 0.17 (0.42) |
| ***$NO_3^-$*** | | | | |
| Exp. 1 - Charcoal vs. Control | 0.287 | 0.076 | **0.019*** | 0.24 (0.49) |
| Exp. 2 - IMOs vs. Control | 0.774 | 0.261 | 0.072 | 0.00 (0.00) |
| Exp. 1 Control vs. Exp. 2 Control | 0.989 | **0.060*** | 0.254 | 0.35 (0.59) |
| ***pH*** | | | | |
| Exp. 1 - Charcoal vs. Control | 0.509 | 0.234 | 0.744 | 0.38 (0.62) |
| Exp. 2 - IMOs vs. Control | 0.361 | **0.007*** | **0.012*** | 0.00 (0.00) |
| Exp. 1 Control vs. Exp. 2 Control | 0.252 | **0.001*** | **0.018*** | 0.16 (0.40) |
| ***Bacteria Shannon diversity*** | | | | |
| Exp. 1 - Charcoal vs. Control | **0.006*** | 0.054 | 0.427 | 0.00 (0.00) |
| Exp. 2 - IMOs vs. Control | 0.205 | **< 0.001*** | 0.450 | 0.32 (0.57) |
| Exp. 1 Control vs. Exp. 2 Control | **0.011*** | **0.009*** | 0.148 | 0.30 (0.55) |
| ***Fungi Shannon diversity*** | | | | |
| Exp. 1 - Charcoal vs. Control | 0.480 | **0.011*** | **0.070*** | 0.04 (0.19) |
| Exp. 2 - IMOs vs. Control | 0.357 | 0.183 | 0.546 | 0.10 (0.31) |
| Exp. 1 Control vs. Exp. 2 Control | **0.014*** | 0.950 | 0.412 | 0.04 (0.20) |

*Models were fitted separately for each of the two bokashi development experiments, and for comparing the controls across the two experiments.

**Table S2.** Relative ASV abundances (%) for bacterial data for each sample used in the bokashi study. Associated metadata for each sample are provided.

**See separate supplementary .xlsx file**

**Table S3.** Relative ASV abundances (%) for fungal data for each sample used in the bokashi study. Associated metadata for each sample are provided.

**See separate supplementary .xlsx file**

**Table S4.** PERMANOVA results of 16S and ITS datasets from bokashi maturation experiments using the Bray-Curtis dissimilarity matrix. Variance explained by each factor.

| **16S** | | |
|---|---|---|
| Model (1) | R2 | p-adjusted |
| **Exp 1. Charcoal vs. Control (16S)** | | |
| Treatment | 0.042 | < 0.001* |
| Time | 0.281 | < 0.001* |
| Treatment x Time | 0.026 | 0.139 |
| **Exp. 2 - IMOs vs. Control (16S)** | | |
| Treatment | 0.041 | < 0.001* |
| Time | 0.365 | < 0.001* |
| Treatment x Time | 0.816 | 0.5894 |
| **Exp. 1 Control vs. Exp. 2 Control** | | |
| Treatment | 0.18 | < 0.001* |
| Time | 0.249 | < 0.001* |
| Treatment x Time | 0.472 | 0.012 |
| **ITS** | | |
| **Exp 1. Charcoal vs. Control (ITS)** | | |
| Treatment | 0.075 | 0.002* |
| Time | 0.344 | 0.002* |
| Treatment x Time | 0.026 | 0.3756 |
| **Exp. 2 - IMOs vs. Control (ITS)** | | |
| Treatment | 0.062 | < 0.001* |
| Time | 0.314 | < 0.001* |
| Treatment x Time | 0.037 | 0.294 |
| **Exp. 1 Control vs. Exp. 2 Control (ITS)** | | |
| Treatment | 0.074 | < 0.001* |
| Time | 0.385 | < 0.001* |
| Treatment x Time | 0.449 | 0.044 |

(1) Models were fitted separately for each of the two bokashi development experiments, and for comparing the controls across the two experiments.

*All p-values are adjusted for multiple tests using Benjamini-Hochberg method.
